# Scanned optogenetic control of mammalian somatosensory input to map input-specific behavioral outputs

**DOI:** 10.1101/2020.08.10.244046

**Authors:** Ara Schorscher-Petcu, Flóra Takács, Liam E. Browne

## Abstract

Somatosensory stimuli guide and shape behavior, from immediate protective reflexes to longer-term learning and high-order processes related to pain and touch. However, somatosensory inputs are challenging to control in awake mammals due to the diversity and nature of contact stimuli. Application of cutaneous stimuli is currently limited to relatively imprecise methods as well as subjective behavioral measures. The strategy we present here overcomes these difficulties by achieving spatiotemporally precise, remote and dynamic optogenetic stimulation of skin by projecting light to a small defined area in freely-behaving mice. We mapped behavioral responses to specific nociceptive inputs and revealed a sparse code for stimulus intensity: using the first action potential, the number of activated nociceptors governs the timing and magnitude of rapid protective pain-related behavior. The strategy can be used to define specific behavioral repertoires, examine the timing and nature of reflexes, and dissect sensory, motor, cognitive and motivational processes guiding behavior.

## Introduction

The survival of an organism depends on its ability to detect and respond appropriately to its environment. Afferent neurons innervating the skin provide sensory information to guide and refine behaviour (Seymour, 2019; Zimmerman et al., 2014). Cutaneous stimuli are used historically to study a wide range of neurobiological mechanisms since neurons densely innervating skin function to provide diverse information as the body interfaces with its immediate environment. These afferents maintain the integrity of the body by recruiting rapid sensorimotor responses, optimize movement through feedback loops, provide teaching signals that drive learning, and update internal models of the environment through higher-order perceptual and cognitive processes (Barik et al., 2018; Brecht, 2017; Corder et al., 2019; de Haan & Dijkerman, 2020; Haggard et al., 2013; Huang et al., 2019; Petersen, 2019; Seymour, 2019). Damaging stimuli, for example, evoke rapid motor responses to minimize immediate harm and generate pain that motivates longer-term behavioral changes.

Compared to visual, olfactory and auditory stimuli, somatosensory inputs are challenging to deliver in awake unrestrained mammals. This is due to the nature of stimuli that require contact and the diversity of stimulus features encoded by afferents that innervate skin. Cutaneous afferent neurons are functionally and genetically heterogeneous, displaying differential tuning, spike thresholds, adaptation rates and conduction velocities (Abraira & Ginty, 2013; Dubin & Patapoutian, 2010; Gatto et al., 2019; Haring et al., 2018). The arborization of their peripheral terminals can delineate spatial and temporal dimensions of the stimulus (Pruszynski & Johansson, 2014), particularly once many inputs are integrated by the central nervous system (Prescott et al., 2014). Cutaneous stimulation in freely moving mice often requires the experimenter to manually touch or approach the skin. This results in inaccurate timing, duration and localization of stimuli. The close proximity of the experimenter can cause observer-induced changes in animal behavior (Sorge et al., 2014). Stimuli also activate a mixture of sensory neuron populations. For example, intense stimuli can co-activate fast-conducting low-threshold afferents that encode innocuous stimuli simultaneously with more slowly-conducting high-threshold afferents (Wang et al., 2018). The latter are nociceptors, that trigger fast protective behaviors and pain. Consequently, mixed cutaneous inputs recruit cells, circuits and behaviors that are not specific to the neural mechanism under study. A way to control genetically-defined afferent populations is to introduce opsins into these afferents and optogenetically stimulate them through the skin (Abdo et al., 2019; Arcourt et al., 2017; Barik et al., 2018; Beaudry et al., 2017; Browne et al., 2017; Daou et al., 2013; Iyer et al., 2014). However, these methods in their current format do not fully exploit the properties of light.

The behaviors that are evoked by cutaneous stimuli are also typically measured with limited and often subjective means. Manual scoring introduces unnecessary experimenter bias and omits key features of behavior. Behavioral assays have traditionally focused on a snapshot of the stimulated body part rather than dynamics of behavior involving the body as a whole (Gatto et al., 2019). Recent advances in machine vision and markerless pose estimation have enabled the dissection of animal behavioral sequences (Mathis et al., 2018; Pereira et al., 2019; Wiltschko et al., 2015). However, these have not been adapted to study behavioral outputs relating to specific cutaneous inputs.

Here we developed an approach to project precise optogenetic stimuli onto the skin of freely-behaving mice (Figure 1A). The strategy elicits time-locked individual action potentials in genetically-targeted afferents innervating a small stimulation field targeted to the skin. Stimuli can be delivered remotely as pre-defined microscale shapes, lines or moving points. The utility of the system was demonstrated by precisely stimulating nociceptors in freely-behaving mice to map behavioral outputs in high-speed. We provide an analysis toolkit that quantifies the millisecond-timescale dynamics of behavioral responses using machine vision methods. We dissect discrete behavioral components of local paw responses, body repositioning and alerting behaviors, and determine how these components relate to the nociceptive input. These data reveal a fundamental neural coding strategy employed by nociceptors to rapidly encode stimulus intensity.

**Figure 1.**
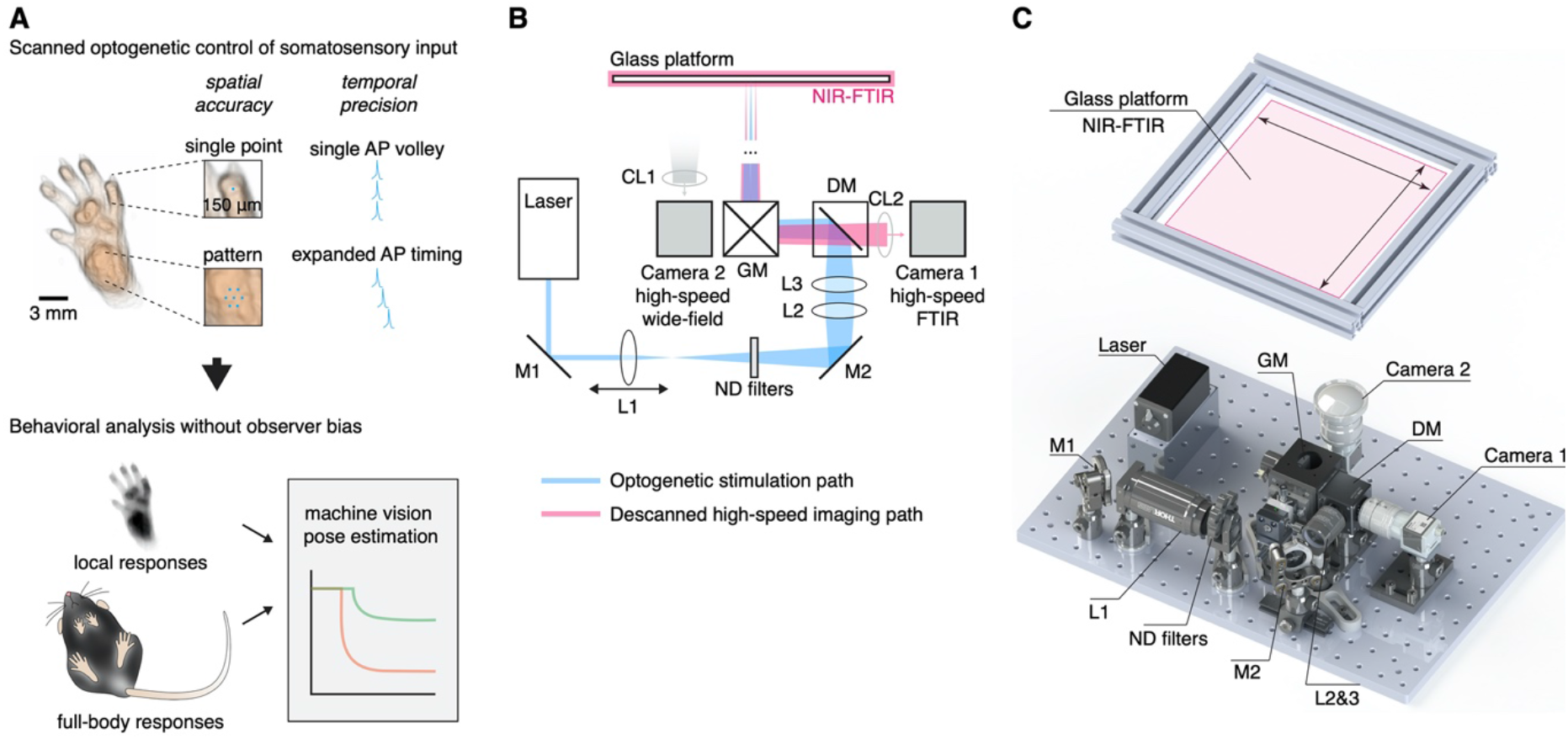
Remote and precise somatosensory input and analysis of behavior. (**A**) The principle, workflow and application of the optical approach. Afferent neurons expressing ChR2 are controlled remotely in freely behaving mice by projecting laser light with sub-millimeter precision to the skin. This enables precise non-contact stimulation with microscale patterns, lines and points using optogenetics. Time-locked triggering of single action potential volleys is achieved through high temporal control of the laser. Behavioral responses can be automatically recorded and analyzed using a combination of machine vision and deep learning methods. (**B**) Schematic of the stimulation laser (in blue) and infrared imaging (in red) paths. Mirrors (M1 and M2) direct the laser beam through a set of lenses (L1-L3), which allow to focus the beam down manually to pre-calibrated sizes. A dichroic mirror (DM) guides the beam into a pair of galvanometer mirrors, which are remotely controlled to enable precise targeting of the beam onto the glass platform. Near-infrared frustrated total internal reflection (NIR-FTIR) signal from the glass platform is descanned through the galvanometers and imaged using a high-speed infrared camera. A second wide-field camera is used to concomitantly record a below-view of the entire glass platform. (**C**) Rendering of the assembled components. A Solidworks assembly is available at https://github.com/browne-lab/throwinglight.

## Results

### Design and assembly of the optical stimulation approach

The design of the optical strategy had eight criteria: (1) that somatosensory stimuli are delivered non-invasively without touching or approaching the mice; (2) localization of stimuli are spatially precise and accurate (<10 μm); (3) freely moving mice can be targeted anywhere within a relatively large (400 cm^2^) arena; (4) stimuli can be controlled with a computer interface from outside the behavior room; (5) stimulation patterns, lines and points are generated by rapidly scanning the stimuli between pre-defined locations; (6) stimulation size can be controlled down to ≥150 μm diameter; (7) stimuli are temporally precise to control individual action potentials using sub-millisecond time-locked pulses; and (8) behavioral responses are recorded at high-speed at the stimulated site and across the whole body simultaneously. An optical system was assembled to meet these criteria (Figure 1B and C).

The stimulation path uses two mirror galvanometers to remotely target the laser stimulation to any location on a large glass stimulation floor. A series of lenses expands the beam and then focuses it down to 0.018 mm^2^ (150 μm beam diameter) at the surface of this floor. This was defocused to provide a range of calibrated stimulation spot sizes up to 2.307 mm^2^, with separable increments that were stable over long periods of time (Figure 1 – figure supplement 1A). The optical power density could be kept equal between these different stimulation spot sizes. The glass floor was far (400 mm) from the galvanometers, resulting in a maximum focal length variability of <1.5% (see Materials and methods). This design yielded a spatial targeting resolution of 6.2 μm while minimizing variability in laser stimulation spot sizes across the large stimulation plane (coefficient of variation ≤0.1, Figure 1 – figure supplement 1B). The beam ellipticity was 74.3 ± 14.3% (median ± MAD, 36–99% range) for all spot sizes. The optical power was uniform across the stimulation plane as expected (Figure 1 – figure supplement 1C). The galvanometers allow rapid small angle step responses to scan the laser beam between adjacent positions and shape stimulation patterns using brief laser pulses (diode laser rise and fall time: 2.5 ns). Custom software (see Materials and methods) was developed to remotely control the laser stimulation position, trigger laser pulses, synchronize galvanometer jumps and trigger the camera acquisition (Figure 1 – figure supplement 2).

The camera acquisition path was used to target the location of the laser stimulation pulse(s); the path was descanned through the galvanometers so that the alignment between the laser and camera is fixed (Figure 1B). High signal-to-noise recordings were obtained using near-infrared frustrated total internal reflection (NIR-FTIR) in the glass stimulation floor (Roberson, D. P. et al., manuscript submitted). If a medium (skin, hair, tail etc.) is within a few hundred microns of the glass it causes reflection of the evanescent wave and this signal decreases non-linearly with distance from the glass such that very minor movements of the paw can be detected. The acquisition camera acquired the NIR-FTIR signal in high-speed (up to 1,000 frames/s) with a pixel size of 110 μm. A second camera was used to record the entire arena and capture behaviors involving the whole body before and after stimulation. Offline quantification was carried out using custom analysis code combined with recently developed markerless tracking tools (Mathis et al., 2018).

### Mapping high-speed local responses to nociceptive input

To validate the strategy, we crossed TRPV1-Cre mice and Cre-dependent ChR2-tdTomato mice, to obtain a line in which ChR2 is selectively expressed in a broad-class of nociceptors innervating glabrous skin (Browne et al., 2017). These TRPV1^Cre^::ChR2 mice were allowed to freely explore individual chambers placed on the stimulation plane. When mice were idle (still and awake), a time-locked laser pulse was targeted to the hind paw. Stimuli could be controlled remotely from outside the behavior room. We recorded paw withdrawal dynamics with millisecond resolution. For example, a single 1 ms laser pulse (stimulation spot size S_6_, 0.577 mm^2^) initiated a behavioral response at 29 ms, progressing to complete removal of the hind paw from the glass floor just 5 ms later (Figure 2A, Figure 2 - video 1). Motion energy, individual pixel latencies, and response dynamics could be extracted from these high-speed recordings (Figure 2B and C).

**Figure 2.**
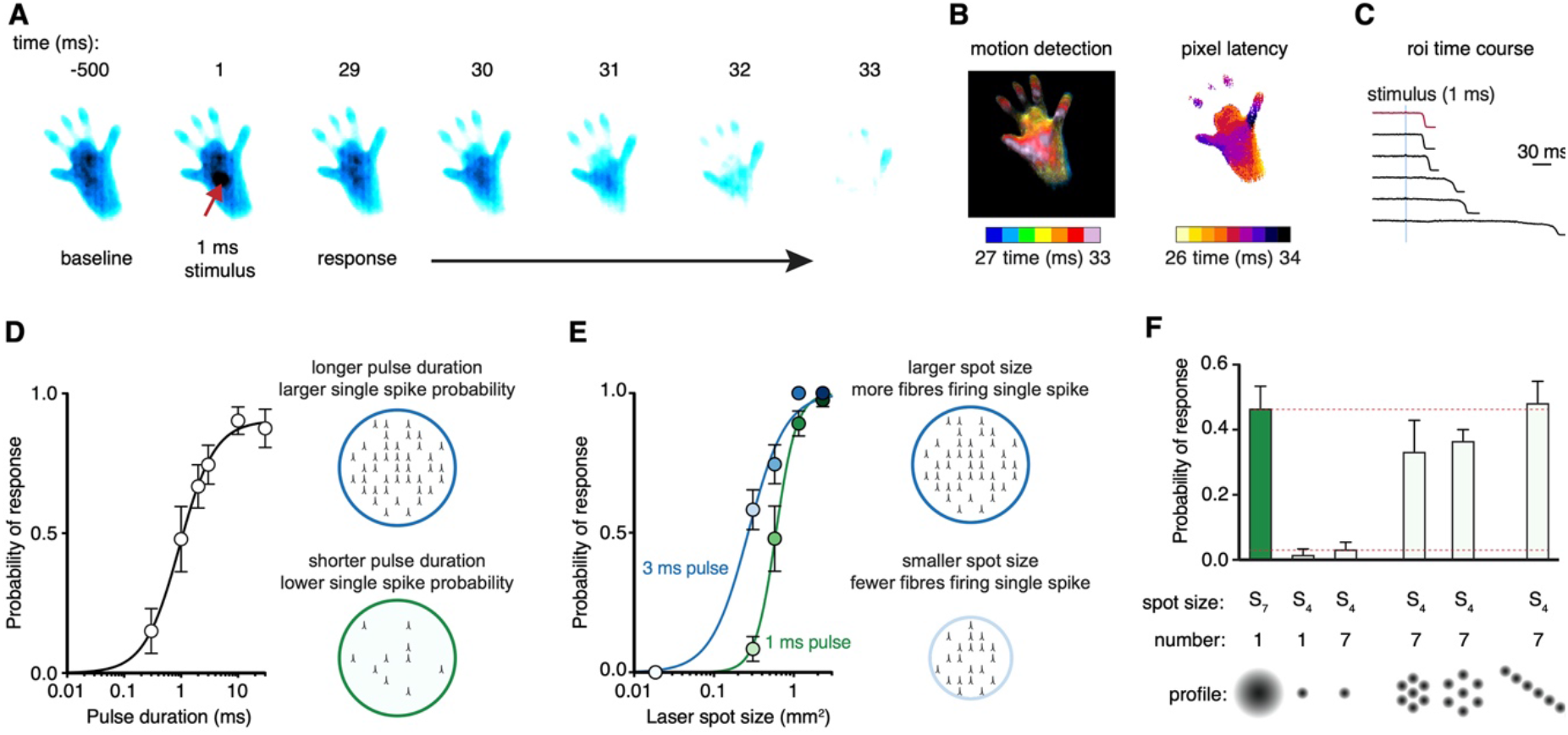
Scanned optogenetic stimuli reveal fast coding of local response probability. (**A**) Millisecond-timescale changes in hind paw NIR-FTIR signal in response to a single 1 ms laser pulse (laser spot size S_6_ = 0.577 mm^2^) recorded at 1,000 frames/s. (**B**) Motion energy analysis (left) and response latencies calculated for each pixel (right) for the same trial as in **A**. (**C**) Example traces of the NIR-FTIR signal time course as measured within a circular region of interest centered on the stimulation site. Six traces from two animals are depicted (1 ms pulse, spot size S_6_ = 0.577 mm^2^). The red trace corresponds to the example trial illustrated in **A**and **B**. (**D**) Paw response probability increases as a function of laser pulse duration when stimulation size is constant (spot size S_6_ = 0.577 mm^2^; 37–42 trials for each pulse duration from n = 8 mice, mean probability ± SEM). (**E**) Paw response probability increases as a function of laser stimulation spot size when pulse duration is constant. Data are 34–45 trials for each spot size per pulse duration from n = 7-8 mice, shown as mean probability ± SEM. (**F**) Stimulation patterning shows that the absolute size, rather than the geometric shape, of the nociceptive stimulus determines the withdrawal probability (Friedman’s non-parametric test for within subject repeated measures S(5) = 22.35, p = 0.0004). Paw response probabilities in response to a single large laser spot (S_7_ = 1.15 mm^2^), a single small spot (S_4_ = 0.176 mm^2^; p = 0.018 compared to S_7_ and p = 0.013 compared to the line pattern), a 10 ms train of seven small 1 ms spots targeting the same site (p = 0.039, compared to S_7_ and p = 0.030 compared to the line pattern) or spatially translated to produce different patterns. Note that the cumulative area of the seven small spots approximates the area of the large spot. Data shown as mean probability ± SEM are from n = 6 mice, with each 6-10 trials per pattern.

We probed multiple sites across the plantar surface and digits and found that the hind paw heel gave the most robust responses (Figure 2 – figure supplement 1). This region was targeted in all subsequent experiments. Littermates that did not express the Cre recombinase allele confirmed that the laser stimulation did not produce non-specific responses. These mice did not show any behavioral responses, even with the largest stimuli (spot size S_8_, 30 ms pulse, Figure 2 – figure supplement 2). We next provide some examples of the utility of the strategy by providing insights into how afferent neurons encode noxious stimuli and generate protective behaviors.

### Precise stimulation reveals sparse coding of response probability

Fast protective withdrawal behaviors can be triggered by the first action potential arriving at the spinal cord from cutaneous nociceptors. A brief optogenetic stimulus generates just a single action potential in each nociceptor activated (Browne et al., 2017). This is due to the rapid closing rate of ChR2 relative to the longer minimal interspike interval of nociceptors. The same transient optogenetic stimulus (Browne et al., 2017), or a pinprick stimulus (Arcourt et al., 2017), initiates behavior before a second action potential would have time to arrive at the spinal cord. That the first action potential can drive protective behaviors places constraints on how stimulus intensity can be encoded, suggesting that the total population of nociceptors firing a single action potential can provide information as a Boolean array. The consequences of this have not been investigated previously as precise control of specific nociceptive input had not been possible. We predicted that the total number of nociceptors firing a single action potential determines features of the behavioral response.

Varying the pulse duration with nanosecond precision influences the probability of each nociceptor generating a single action potential within the stimulation site. A pulse as short as 300 μs elicited behavioral responses but with relatively low probability (Figure 2D). This probability increased with pulse duration until it approached unity, closely matching the on-kinetics of the ChR2 used (*τ* =1.9 ms (Lin, 2011)). We next controlled the spatial, rather than temporal, properties of the stimulation in two further experiments. Firstly, we find that the total area of stimulated skin determines the behavioral response probability, such that the larger the nociceptive input the larger the response probability (Figure 2E). Secondly, we generated different stimulation patterns. We find that sub-threshold stimulations are additive (Figure 2F). Specifically, seven spatially displaced small subthreshold stimulations could reproduce the response probability of a single large stimulation that was approximately seven times their size. This could not be achieved by repeated application of the small stimulations to the same site (Figure 2F).

### Sparse coding of local response latency and magnitude

We examined the response dynamics of the stimulated hind paw. Time-locked stimulation of the hind paw (Figure 3A) resulted in responses that were analyzed using a hierarchical bootstrap estimate of the median (see Materials and methods). The nociceptive input size influenced the behavioral response latency: for example, a 3 ms pulse resulted in bootstrap response latencies of 27 ± 1 ms, 30 ± 2 ms, 33 ± 5 ms and 112 ± 46 ms were determined for spot sizes S8, S7, S6 and S5, respectively (Figure 3B). The shorter latencies are consistent with medium-conduction velocity Aδ-fibres (Arcourt et al., 2017; Browne et al., 2017). The rank order of response latencies follows the nociceptive input size for both pulse durations, and they fit well with log-log regressions (3 ms pulse *R^2^* = 0.87, 1 ms pulse *R^2^* = 0.90). Once a hind limb motor response was initiated it developed rapidly, lifting from the glass with bootstrap rise times that show the vigor of the motor response was also dependent on nociceptive input magnitude (Figure 3C). These responses, in >65% of cases, proceeded to full withdrawal. However, in a fraction of trials the paw moved but did not withdraw (Figure 3D). Notably, such responses are not detected by eye, highlighting the sensitivity of the acquisition system. Even the smallest of nociceptive inputs still produced a large fraction of full withdrawal responses, despite decreases in response probability (Figure 3E). The fraction of full withdrawal responses increased with the size of nociceptive input. The onset latency of both full and partial responses decreased as nociceptive input increased (Figure 3F).

**Figure 3.**
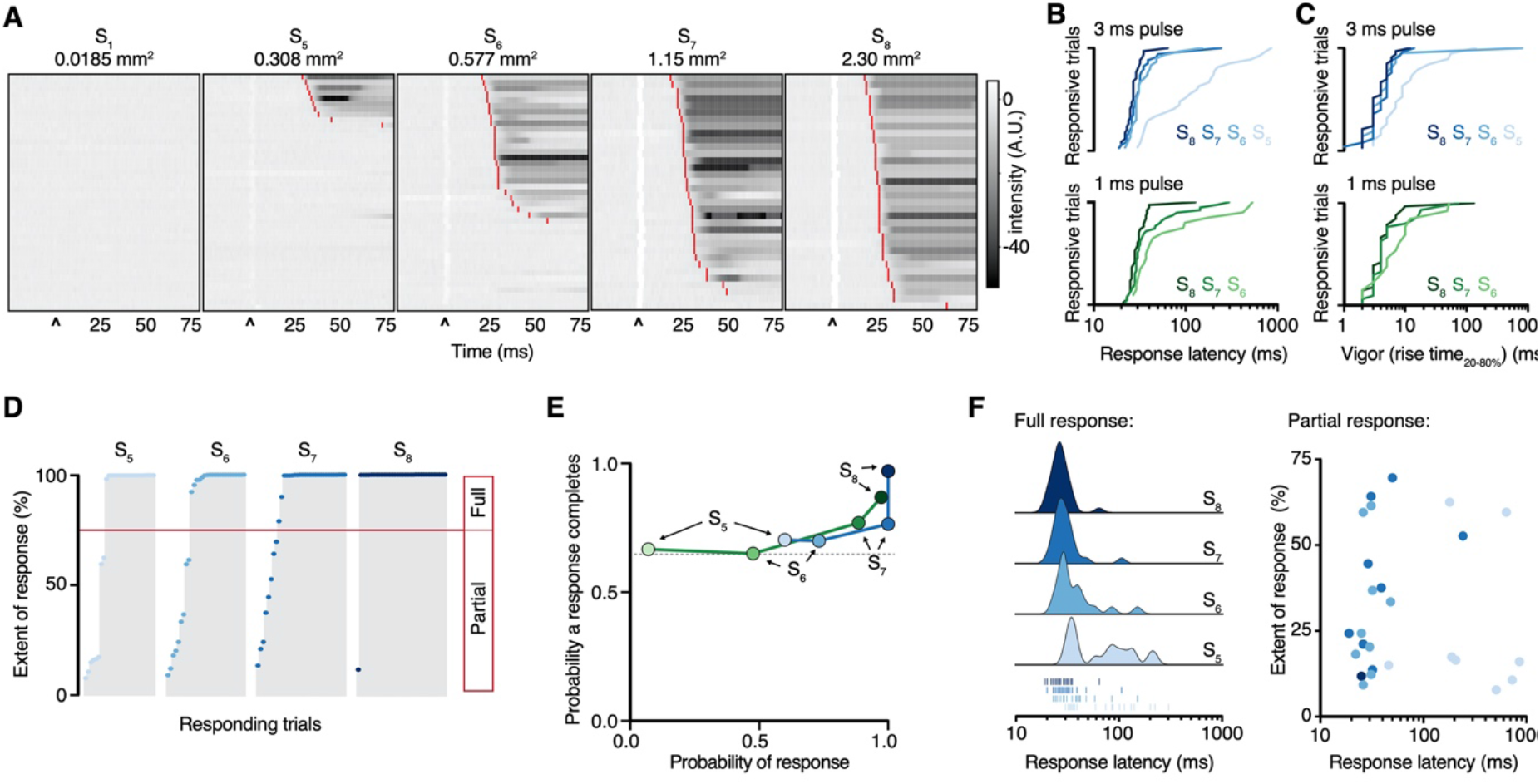
Paw response latency and magnitude is influenced by a sparse code. (**A**) Raster plots of hind paw dynamics for five different 3 ms laser stimulation spot sizes sorted by response latency. The paw response latency is indicated in red. (**B**) Paw response latencies to trials with single 3 ms (blue, left) and 1 ms (green, right) stimulations at different spot sizes, sorted by latency. (**C**) Response vigor (hind paw rise time, 20-80%) to single 3 ms (blue, left) or 1 ms (green, right) pulses with a range of stimulation spot sizes. Bootstrap rise times to a 3 ms pulse were 4 ± 1 ms, 4 ± 1 ms, 4 ± 1 ms and 9 ± 5 ms for spot sizes S8, S7, S6 and S5, respectively, and to a 1 ms pulse were 4 ± 1 ms, 5 ± 2 ms and 6 ± 3 ms for spot sizes S8, S7 and S6, respectively. (**D**) Extent of response (%NIR-FTIR signal decrease). The threshold for a full response and partial response is 75% of baseline signal (red line). (**E**) The probability of responses to reach completion (full response) as a function of the probability of response for four stimulation spot sizes and two pulse durations (green 1 ms; blue 3 ms). (**F**) Response latency distributions for trials that reach completion (full response) shown with Gaussian kernel density estimation of data (left). Rug plot inset representing individual response latencies for each color-coded spot size. No correlation was observed between response latency and extent for partial responses when stimulation duration was 3 ms. Data is from 8 mice each with six trials (48 trial total). After automated quality control the trial numbers for the 1 ms stimulus duration were: 43 trials from 8 mice for spot size S5 (3 responses: 2 full and 1 partial); 42 trials from 8 mice for spot size S6 (20 responses: 13 full and 7 partial); 44 trials from 8 mice for spot size S7 (39 responses: 30 full and 9 partial); and 39 trials from 7 mice for spot size S8 (38 responses: 33 full and 5 partial). Similarly, the trial numbers for the 3 ms stimulus duration were: 44 trials from 8 mice for spot size S5 (27 responses: 19 full and 8 partial); 41 trials from 8 mice for spot size S6 (30 responses: 21 full and 9 partial); 34 trials from 8 mice for spot size S7 (34 responses: 26 full and 8 partial); and 34 trials from 7 mice for spot size S8 (34 responses: 33 full and 1 partial).

### Full-body behavioral responses to remote and precise nociceptive input

Pain-related responses are not limited to the affected limb but involve simultaneous movement of other parts of the body (Blivis et al., 2017; Browne et al., 2017). These non-local behaviors theoretically serve several protective purposes: to investigate and identify the potential source of danger, move the entire body away from this danger, attend to the affected area of the body (Huang et al., 2019) and to maintain balance (Sherrington, 1910). Full-body movements were quantified as motion energy (Figure 4A) and high-speed recordings show this initiated with a bootstrap mean response latency of 30 ± 1 ms and the first movement bout had a bootstrap mean duration of 136 ± 14 ms (80 trials from 10 mice) (Figure 4 – figure supplement 1). The magnitude of full-body movement increased with the stimulation spot size (Figure 4B). Bootstrap peak motion energy had a lognormal relationship with nociceptive input size (*R^2^* = 0.99). This indicates global behaviors are also proportional to the number of nociceptors that fire a single action potential (Figure 4B).

**Figure 4.**
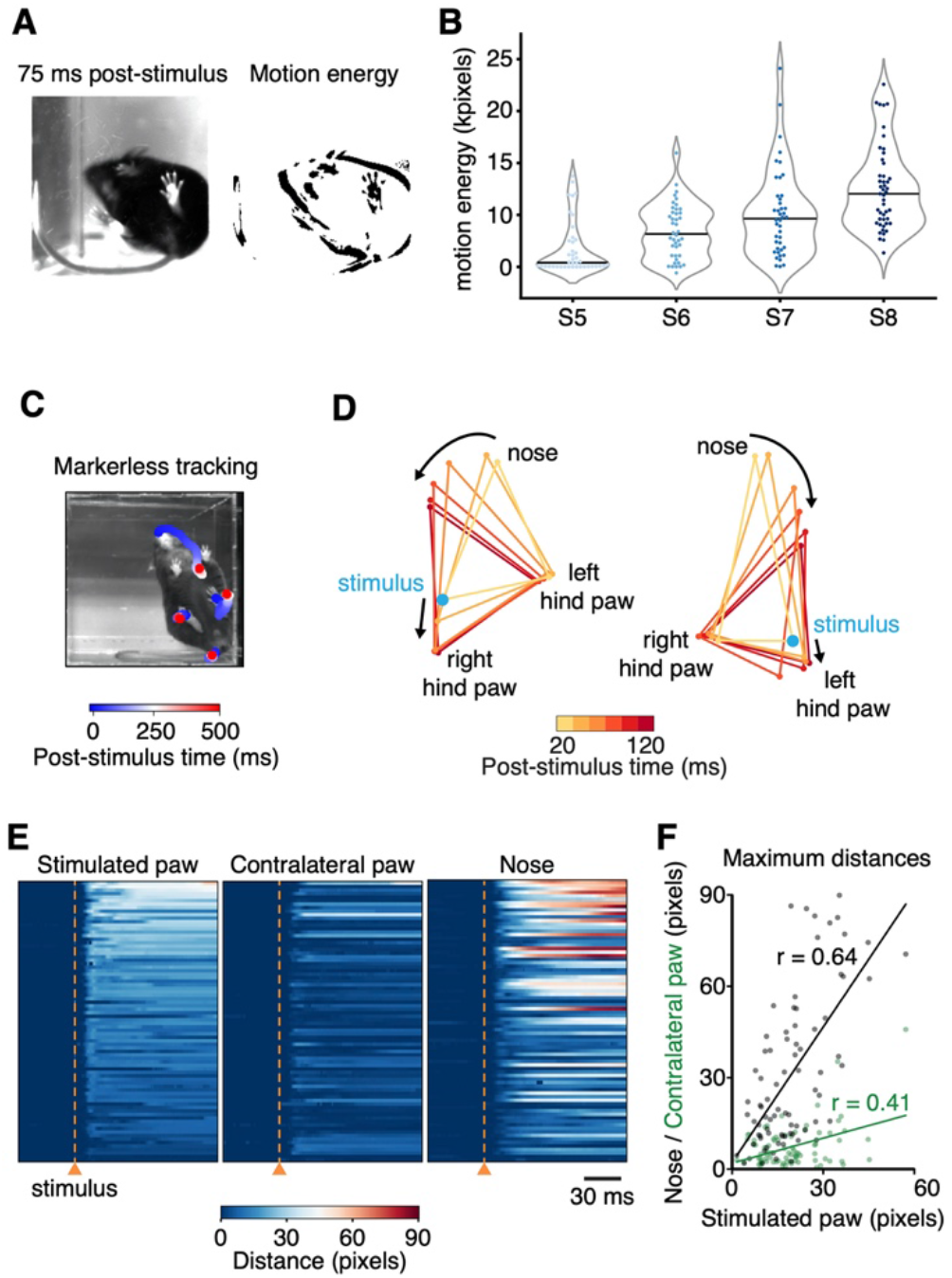
Dissection of full-body behavioral response repertoire to precise nociceptive input. (**A**) Left: Example image from the below-view camera, recording whole-body behavior with 40 frames/s, 75 ms after stimulus delivery (3 ms pulse, spot size S_6_ = 0.577 mm^2^). Right: Visual representation of motion energy calculated 75 ms after the stimulus. (**B**) Motion energy increases with larger spot sizes when pulse duration is kept constant at 3 ms. Violin plots with 41 to 47 trials per spot size from 8 mice. Individual trials are shown, along with the associated median in black. (**C**) Example spatiotemporal structure of a noxious stimulus response superimposed on the baseline image taken immediately before stimulus. The color indicates the timing of nose and hind paw trajectories. In this example, the left side of the mouse was stimulated. (**D**) Example graphical representation showing the sequence of postural adjustment following nociceptive stimulus. Left: the right hind paw of the mouse was stimulated. Right: the left hind paw of the mouse was stimulated. (**E**) Summary raster plots of the distances that each tracked body part moves in n = 80 trials (from 10 mice). All raster plots are sorted by maximum distances achieved by the stimulated paw within 300 ms of the stimulation. (**F**) Correlations of maximum distances traveled within 300 ms of stimulation by the nose (black) and contralateral paw (green) and the stimulated hind paw for all trials shown in **E**.

### Nociceptor sparse coding triggers coordinated postural adjustments

Markerless tracking of individual body parts can reveal the coordination of behavioral responses (Figure 4C). We tracked 18 sites across the body of the mouse at high-speed (400 frames/s) and quantified behavioral response dynamics, extent and coordination to an intense hind paw stimulus (S8, 2.307 mm^2^, 10 ms pulse) for three of these sites (Figure 4D and E). Bootstrap response latencies show fast outward movement of the stimulated paw (29 ± 1 ms) and contralateral paw (34 ± 4 ms), and concomitant initiation of head orientation (33 ± 2 ms, 80 trials from 10 mice). With this intense stimulus, only in 6% of trials did the hind paws or single body parts move alone, although the strength of the head orientation varied between trials (Figure 4E). Quantification of the displacement of each body part relative to its baseline position reveals a positive correlation between distances traveled by the nose and the stimulated paw (Pearson’s r = 0.64, Figure 4F, n = 80 trials from 10 mice). The presence of head orientation suggests that a brief nociceptive input can rapidly generate a coordinated spatially organized behavioral response that aims to gather sensory information about the stimulus or its consequences, and potentially provides coping strategies. Protective behaviors can be statistically categorized (Abdus-Saboor et al., 2019). We have shown that the analysis can easily be customized to incorporate computational tools that facilitate quantification and reveal insights into complex behavioral responses.

## Discussion

We describe a strategy for remote, precise, dynamic somatosensory input and behavioral mapping in awake unrestrained mice. The approach can remotely deliver spatiotemporally accurate optogenetic stimuli to the skin with pre-defined size, geometry, duration, timing and location, while simultaneously monitoring behavior in the millisecond timescale. Action potentials can be generated asynchronously by altering the sub-millisecond timings of each light pulse in a patterned stimulus. Microscale optogenetic stimulation can be used to simulate textures, edges and moving points on the skin. Responses to these precisely defined points and patterns can be mapped using machine vision approaches. The design is modular, for example additional lasers for multicolor optogenetic control or naturalistic infrared stimuli can be added and complementary machine vision analysis approaches readily implemented.

We validated the system in a transgenic mouse line providing optical control of a broad class of nociceptors. Advances in transcriptional profiling have identified a vast array of genetically-defined primary afferent neuron populations involved in specific aspects of temperature, mechanical and itch sensation (Usoskin et al., 2015). Selective activation of these populations is expected to recruit a specific combination of downstream cells and circuits depending on their function. For example, nociceptive input generates immediate sensorimotor responses and also pain that acts as a teaching signal. This strategy can be thus combined with techniques to modify genes, manipulate cells and neural circuits, and record neural activity in freely behaving mice to probe these mechanisms (Boyden et al., 2005; Kim et al., 2017). We provide approaches to map behavioral responses to defined afferent inputs across the spectrum of somatosensory modalities (Browne et al., 2017; Huang et al., 2019).

We find that the probabilistic recruitment of nociceptors can serve as a code for noxious stimulus intensity. This determines the behavioral response probability, latency and magnitude. In contrast to firing-rate dependent codes for low-threshold mechanoreceptive afferents (Muniak et al., 2007) and nociceptors (Wang et al., 2018; Yarmolinsky et al., 2016), this code utilizes the total number of first action potentials arriving at the spinal cord, rather than information from trains of action potentials that might delay protective response times. Therefore, it resembles a fast and sparse population code, where nociceptor spikes are summated by spinal neurons and trigger behavior when certain thresholds are exceeded. This neural mechanism is separate from time delays related to temperature changes or mechanical deformation in the skin (Danneman et al., 1994). The delay to withdraw from a hot surface, for example, is not simply the time it takes to heat the skin but is determined by the total number of first action potentials encoding the stimulus. The intensity, size and location of a stimulus can be conveyed rapidly by this neural code. Relative arrival times of the first action potentials might also contribute to the code, as observed in the visual system (Gollisch & Meister, 2008), and subsequent action potentials could enable multiplexing (Lankarany et al., 2019). We use a broad-class nociceptor line and it is possible that its subpopulations exploit a diversity of coding strategies. This optical approach can reveal how such subpopulations and their specific downstream circuits guide behavior.

In summary, we have developed a strategy to precisely control afferents in the skin without touching or approaching them, by projecting light to optogenetically generate somatosensory input in patterns, lines or points. This is carried out non-invasively in awake unrestrained mammals in a way that is remote yet precise. Remote control of temporally and spatially precise input addresses the many limitations of manually applied contact stimuli. The timing, extent, directionality and coordination of resultant millisecond-timescale behavioral responses can be investigated computationally with specific sensory inputs. This provides a way to map behavioral responses, circuits and cells recruited by defined afferent inputs and to dissect the neural basis of processes associated with pain and touch. This strategy thus enables the investigation of sensorimotor, perceptual, cognitive and motivational processes that guide and shape behavior in health and disease.

## Materials and methods

### Optical system design, components and assembly

Optical elements, optomechanical components, mirror galvanometers, the diode laser, LEDs, controllers, machine vision cameras, and structural parts for the optical platform are listed in Figure 1 – table 1. These components were assembled on an aluminum breadboard as shown in the Solidworks rendering in Figure 1C. The laser was aligned to the center of all lenses and exiting the midpoint of the mirror galvanometer housing aperture when the mirrors were set to the center of their working range. A series of lenses (L1-L3) expanded the beam before focusing it on to the glass stimulation plane, on which mice are placed during experiments. The glass stimulation platform was constructed of 5 mm thick borosilicate glass framed by aluminum extrusions. Near-infrared frustrated total internal reflection (NIR-FTIR) was achieved by embedding an infrared LED ribbon inside the aluminum frame adjacent to the glass edges (Roberson, D. P. et al., manuscript submitted). The non-rotating L1 lens housing was calibrated to obtain eight defined laser spot sizes, ranging from 0.0185 mm^2^ to 2.307 mm^2^, by translating this lens along the beam path at set points to defocus the laser spot at the 200 mm x 200 mm stimulation plane. To ensure a relative flat field in the stimulation plane, the galvanometer housing aperture was placed at a distance of 400 mm from its center. In this configuration, the corners of the stimulation plane were at a distance of 424 mm from the galvanometer housing aperture and variability of the focal length was below 1.5%.

Optical power density was kept constant by altering the laser power according to the laser spot area. Neutral density (ND) filters were used so that the power at the laser aperture was above a minimum working value (≥8 mW) and to minimize potential changes in the beam profile at the stimulation plane. The laser and mirror galvanometers were controlled through a multifunction DAQ (National Instruments, USB-6211) using custom software written in LabVIEW. The software displays the NIR-FTIR camera feed, whose path through the mirror galvanometers is shared with the laser beam, so that they are always in alignment with one another. Computationally adjusting mirror galvanometer angles causes identical shifts in both the descanned NIR-FTIR image field of view and intended laser stimulation site, so that the laser can be targeted to user-identified locations. Shaped stimulation patterns were achieved by programmatically scaling the mirror galvanometer angles to the glass stimulation plane using a calibration grid array (Thorlabs, R1L3S3P). The timings of laser pulse trains were synchronized with the mirror galvanometers to computationally implement predefined shapes and lines using small angle steps that could be as short as 300 μs. The custom software also synchronized image acquisition from the two cameras, so that time-locked high-speed local paw responses were recorded (camera 1: 160 pixels x 160 pixels, 250-1,000 frames/s depending on the experiment). Time-locked global full-body responses were recorded above video-frame rate (camera 2: 664 pixels x 660 pixels, 40 frames/s) or at high-speed (camera 2: 560 pixels x 540 pixels, 400 frames/s) across the entire stimulation platform.

### Technical calibration and characterization of the optical system

To calibrate the L1 lens housing and ensure consistency of laser spot sizes across the glass stimulation platform we designed a 13.90 ± 0.05 mm thick aluminum alignment mask. This flat aluminum mask was used to replace the glass stimulation platform and was combined with custom acrylic plates that align the aperture of a rotating scanning-slit optical beam profiler (Thorlabs, BP209-VIS/M) to nine defined coordinates at different locations covering the stimulation plane. The laser power was set to a value that approximates powers used in behavioral experiments (40 mW). The laser power was then attenuated with an ND filter to match the operating range of the beam profiler. Using Thorlabs Beam Software, Gaussian fits were used to determine x-axis and y-axis 1/e^2^ diameters and ellipticities for each laser spot size over three replicates at all nine coordinates. The averages of replicates were used to calculate the area of the eight different laser spot sizes that were measured in each of the nine coordinates (Figure 1 – figure supplement 1A) and then fitted with a two-dimensional polynomial equation in MATLAB to create heatmaps (Figure 1 – figure supplement 1 B).

The average values over the nine coordinates were defined for each laser spot size: S_1_ = 0.0185 mm^2^, S_2_ = 0.0416 mm^2^, S_3_ = 0.0898 mm^2^, S_4_ = 0.176 mm^2^, S_5_ = 0.308 mm^2^, S_6_ = 0.577 mm^2^, S_7_ = 1.155 mm^2^, S_8_ = 2.307 mm^2^. These measurements were repeated six months after extensive use of the optical system to ensure stability over time (Figure 1 – figure supplement 1A). In addition, the uniformity of laser power was assessed by measuring optical power at five positions of the experimental platform with a power meter (Thorlabs, PM100D) (Figure 1 – figure supplement 1C).

### Experimental animals

Experiments were performed using mice on a C57BL/6j background. Targeted expression of ChR2-tdTomato in broad-class cutaneous nociceptors was achieved by breeding mice homozygous for Cre-dependent ChR2(H134R)-tdTomato at the ROSA26 locus (RRID: IMSR_JAX:012567, Ai27D, ChR2-tdTomato) (Madisen et al., 2012) with mice that have Cre recombinase inserted downstream of the *Trpv1* gene in one allele (RRID:IMSR_JAX:017769 JAX 017769, TRPV1^Cre^) (Cavanaugh et al., 2011). Resultant mice were heterozygous for both transgenes and were housed with control littermates that do not encode Cre recombinase but do encode Cre-dependent ChR2-tdTomato. Adult (2–4 months old) male and female mice were used in experiments. Mice were given *ad libitum* access to food and water and were housed in 21°C ± 2°C, 55 % relative humidity and a 12 hr light:12 hr dark cycle. Experiments were typically carried out on a cohort of 4 to 6 mice and spaced by at least one day in the case where the same cohort of mice was used in different experiments. All animal procedures were approved by University College London ethical review committees and conformed to UK Home Office regulations.

### Optogenetic stimulation and resultant behaviors

Prior to the first experimental day, mice underwent two habituation sessions during which each mouse was individually placed in a plexiglass chamber (100 mm x 100 mm, 130 mm tall) on a mesh wire floor for one hour, then on a glass platform for another hour. On the experimental day, mice were again placed on the mesh floor for one hour, then up to six mice were transferred to six enclosures (95 mm x 60 mm, 75 mm tall) positioned on the 200 mm x 200 mm glass stimulation platform. Mice were allowed to settle down and care was taken to stimulate mice that were calm, still and awake in an “idle” state. The laser was remotely targeted to the hind paw glabrous skin using the descanned NIR-FTIR image feed. The laser spot size was manually set using the calibrated L1 housing, while laser power and neutral density filters were used to achieve a power density of 40 mW/mm^2^ regardless of spot size. The software was then employed to trigger a laser pulse of defined duration (between 100 μs and 30 ms) and simultaneously acquire high-speed (1,000, 500 or 250 frames/s depending on experiment) NIR-FTIR recordings of the stimulated paw, as well as a global view of the mice with a second camera (40 frames/s or 400 frames/s) (Figure 1C). Each recording was 1,500 ms in duration, with the laser pulse initiated at 500 ms. The behavioral withdrawal of the stimulated hind paw was also manually recorded by the experimenter. For each stimulation protocol, 6 pulses, 3 on each hind paw, spaced by at least one minute were delivered to eight mice, split into two cohorts.

### Patterned stimulation protocols

Mice were stimulated on the heel of the hind paw with each of the following protocols: (1) a single 1 ms pulse with spot size S_7_ (1.155 mm^2^); (2) a single 1 ms pulse with spot size S_4_ (0.176 mm^2^); (3) seven 1 ms pulses with spot size S_4_, superimposed on the same stimulation site and spaced by 500 μs intervals; (4) seven 1 ms pulses with spot size S_4_, spaced by 500 μs intervals and spatially displacing stimuli with 0.3791 mm jumps such as to draw a small hexagon; (5) seven 1 ms pulses with spot size S_4_, spaced by 500 μs intervals and spatially displacing stimuli with 0.5687 mm jumps such as to draw a hexagon expanded by 50% compared to the previous shape; (6) seven 1 ms pulses with spot size S_4_, spaced by 500 μs intervals and spatially displacing stimuli with 0.3791 mm jumps such as to draw a straight line. The power density of the stimulations was kept constant at 40 mW/mm^2^ as before. Seven mice, split into two cohorts, received ten stimulations per protocol (five on each hind paw) after a baseline epoch of 500 ms. An additional cohort of four littermates carrying a wild-type locus at the Trpv1-Cre allele were stimulated in the same way and served as negative controls. Finally, three TRPV1-Cre::ChR2 mice were stimulated (spot size S_8_, 10 ms pulse duration) with a single pulse adjacent to the hind paw, five times on each side, in order to control for potential off-target effects. The NIR-FTIR signal was recorded at 500 frames/s.

### Global behaviors during optogenetic stimulation

To obtain recordings optimized for markerless tracking with DeepLabCut, a single acrylic chamber (100 mm x 100 mm, 150 mm tall) was centered on the glass stimulation platform of the system. Rapid movements were recorded at 400 frames/s using a below-view camera (FLIR, BFS-U3-04S2M-CS). Two white and two infrared LED panels illuminated the sides of the behavioral chamber in order to optimize lighting for these short exposure times and achieve high contrast images. NIR-FTIR was not used in this configuration. Mice received between 10 and 20 single-shot laser pulse stimulations of 10 ms each, at least 1 minute apart and equally split between right and left hind paw and using spot size S_8_ (2.31 mm^2^). The first 10 trials that exceeded DeepLabCut quality control were used. Each trial consisted of a 500 ms baseline and 4,000 ms after-stimulus recording epoch.

### Automated analysis of optogenetically evoked local withdrawal events

High-speed NIR-FTIR recordings were saved as uncompressed AVI files. A python script was implemented in Fiji to verify the integrity of the high-speed NIR-FTIR recordings and extract average 8-bit intensity values from all frames within a circular region of interest on the stimulation site (60 pixels diameter). This output was then fed into Rstudio to calculate the average intensity and associated standard deviation of the baseline recording (first 500 ms). A hind paw response was defined as a drop of intensity equal to or below the mean of the baseline minus five times its standard deviation. Paw response latency was defined as time between the start of the pulse and the time at which a hind paw response was first detected. For purposes of quality control, only recordings with a baseline NIR-FTIR intensity mean ≥3 and a standard deviation/mean of the baseline ratio ≥23 were retained for analysis. Another criterion was that response latencies are not 10 ms or shorter since this would be too short to be generated by the stimulus itself. Only one trial out of 2369 trials did not meet this criterion (spot size S_6_, 1ms pulse, 8 ms response latency). In addition to this two-step work-flow using Fiji/Python to process AVI files and then Rstudio to analyze the resulting output, alternative code was written in Python 3, which combines both steps and also computes individual pixel latencies and motion energy using NumPy and Pandas packages. A median filter (radius = 2 pixels) was applied to the NIR-FTIR recordings used to create the representative time-series in Figure 2A and Figure 2 – video 1. For raster plots of hind paw response dynamics in Figure 4A, NIR-FTIR intensity values were normalized to the average baseline value. For the patterned stimulation experiments in Figure 2F, trials were analyzed as stated to compute local response probabilities, but an additional rule was introduced to further minimize the risk of false positives. A response required the signal to fall by 20% and exceed a threshold of four times the standard deviation of baseline.

### Automated analysis of full-body protective behavior

Videos of the entire stimulation platform were cropped into individual mouse chambers (200 x 315 pixels) and then analyzed using Rstudio to quantify the amount of full-body movements, including those stemming from the response of the stimulated limb, herein referred to as global behavior (GB). GB was approximated as the binarized motion energy: the summed number of pixels changing by more than five 8-bit values between two subsequent frames (Pixel Change). Briefly, for each pixel_*n*_ (n = 63,000 pixels/frame), the 8-bit value at a given frame (*F_n_*) was subtracted from the corresponding pixel_*n*_ at the previous frame (*F_n-1_*). If the resulting absolute value was ≤5, 0 would be assigned to the pixel. If the absolute resulting value was >5, 1 would be assigned to the pixel. The threshold was chosen to discard background noise from the recording. The pixel binary values were then summed for each frame pair to obtain binarized motion energy. Normalized binarized motion energy was calculated by subtracting each post-stimulus frame binarized motion energy from the average baseline binarized motion energy. As an alternative to this analysis strategy, we have developed code in Python that processes the video files and calculates motion energy. The peak normalized binarized motion energy within a 75 ms time window (first three frame pairs proceeding the stimulus) was determined and only trials displaying a peak response ≥5 standard deviations of the baseline mean were retained for further analysis and plotting. Between 41 and 47 videos from 8 mice were analyzed per spot size.

### Markerless tracking of millisecond-timescale global behaviors

#### DeepLabCut installation

DeepLabCut (version 2.0.1) was installed on a computer (Intel®-Core™-i7-7800X 3.5 GHz CPU, NVIDIA GTX GeForce 1080 Ti GPU, quad-core 64 GB RAM, Windows 10, manufactured by PC Specialist Ltd.) with an Anaconda virtual environment and was coupled to Tensorflow-GPU (v.1.8.0, with CUDA v.9.01 and cUdNN v. 5.4).

#### Data compression

All recordings were automatically cropped with python MoviePy package and compressed with standard compression using the H.264 codec, then saved in mp4 format. This compression method was previously shown to result in robust improvement of processing rate with minimal compromise on detection error.

#### Training the network

DeepLabCut was used with default network and training settings. Pilot stimulation trials were collected for initial training with 1,030,000 iterations from 253 labeled images from 50 videos. The videos were selected to represent the whole range of behavioral responses and conditions (25 videos of males and 25 videos of females from six different recording sessions). Out of the 25 videos, 15 were selected from the most vigorous responses, five were selected from less vigorous responses and five from control mice. Ground truth images were selected manually, aiming to include the most variable images from each video (up to 14 frames per video). 18 body parts were labeled, namely the nose, approximate center of the mouse, two points on each sides of the torso and one point at each side of the neck, the fore paws, distal and proximal points on the hind paw, between the hind limbs, and three points on the tail. While most of these labels were not used in subsequent analysis, labeling more body parts on the image enhanced performance. The resulting network output was visually assessed. Erroneously labeled frames were manually corrected and used to retrain the network while also adding new recordings. Four sequential retraining sessions with 1,030,000 iterations each were conducted adding a total of 109 frames from 38 videos. This resulted in a reduction in the pixel RMSE (root mean square error) from 4.97 down to 2.66 on the test set, which is comparable to human ground truth variability quantified elsewhere.

#### Data processing

Only labels of interest were used for analysis. These were ipsilateral and contralateral hind paws (distal), the tail base and the nose labels. To minimize error, points were removed if: 1) they were labeled with less than 0.95 p-cutoff confidence by DeepLabCut; 2) they jumped at least 10 pixels in one single frame compared to the previous frame; 3) they had not returned on the subsequent frame; and 4) they were from the 5 stimulation frames. Code for data processing was written in Python using the NumPy and Pandas packages. Additional post-hoc quality control was performed on the network output to identify and remove poorly labeled trials. To this end, heat maps of distances between labels were created and inspected for dropped labels and sudden changes in distance. Trials identified in this manner were then manually inspected and removed if more than 10% of labels were missing or more than 10 frames were mislabeled. In total, 4.7% of trials were discarded. Only the first 8 trials for each of the 10 mice that met this video quality control were used in analysis.

#### Automated detection of the stimulated limb

Disabling NIR-FTIR illumination reduces the baseline saturation and thus allowed us to automate stimulated paw detection using pixel saturation from the stimulation laser. To determine which of the left or right paw had been stimulated in a given trial, the number of saturated pixels within a 60 x 60 pixels window close to the hind paw label were compared 7.5 ms prior and 5 ms after stimulus onset.

#### Detection of movement latency of discrete body parts

Movement latencies of hind paws and head (nose) were computed based on significant changes from the baseline position. Baseline positions were calculated as the average x and y values from 10 consecutive frames prior to stimulus onset. A post-stimulus response was considered to be meaningful if the position of the label changed by at least 0.5 pixels (~0.16 mm) compared to baseline and continued moving at a rate of at least 0.5 pixel/frame for the subsequent 10 frames.

### Motion energy calculations in millisecond-timescale global behaviors

GB was analyzed within a 1 ms time frame following stimulation by computing the binarized motion energy relative to a baseline reference frame 5 ms prior to stimulation as described above. Here, the threshold for pixel change was set to seven 8-bit values. The binarized motion energy (sum of pixel binaries) of a given frame was normalized to the total number of pixels within that frame after removing those frames that had been affected by the stimulation laser pulse. The global response latency of movement initiation was determined as the time when binarized motion energy was greater than 10 times the standard deviation at baseline. Termination of movement was determined as the time point when binarized motion energy returned below 10 times standard deviation from baseline following the first movement bout.

### Statistical Analysis

Data was analyzed in Rstudio 1.2.5019, Python 3.6.8, ImageJ/FIJI 2.0.0 and Prism 7 and visualized using Seaborn, Prism 7 and Adobe Illustrator 24.0. In all experiments repeated measurements were taken from multiple mice. Paw responses to patterned stimulation were reported as mean probabilities ± standard error of the mean (SEM) and analyzed using Friedman’s non-parametric test for within-subject repeated measures followed by Dunn’s signed-rank test for multiple comparisons (Figure 2F). In this experiment, one of the seven TRPV1-Cre::ChR2 mice was removed from the data set because it displayed saturating responses to Protocol 3 preventing comparison of values across a dynamic range. Response latencies, response rise times and response durations were computed using a hierarchical bootstrap procedure (Saravanan et al., 2019) modified to acquire bootstrap estimates of the median with balanced resampling. Briefly, mice are sampled with replacement for the number of times that there are mice. For each mouse within this sample its trials were sampled with replacement, but the number of selected trials were balanced, ensuring each mouse contributes equally to the number of trials in the sample. The median was taken for this resampled population and this entire process was repeated 10,000 times. Values provided are the mean bootstrap estimate of the median ± the standard error of this estimate. The median bias was small due to the resampled population size from hierarchically nested data and only moderate distribution skew. Global peak motion energy (Figure 4B) was examined in a similar way, except the mean of resampled populations was used as it represents a better estimator of the population mean. In this case, we report the mean bootstrap estimate of the mean ± the standard error of this estimate. Pearson’s correlation coefficients were determined to compare maximum distances moved from baseline for each body part (Figure 4F). Experimental units and n values are indicated in the figure legends.

## Data and code availability

All components necessary to assemble the optical system are listed in Figure 1 – table 1. A Solidworks assembly, the optical system control and acquisition software and behavioral analysis toolkit are available at https://github.com/browne-lab/throwinglight. The data that support the findings of this study are available from the corresponding author upon reasonable request.

## Author Contributions

L.E.B. conceived and built the optical system and wrote the control and acquisition software. A.S.-P. and L.E.B. designed experiments and wrote the manuscript. A.S.-P., F.T. and L.E.B. carried out experiments, wrote code, analyzed data and interpreted results.

## Acknowledgments

We are grateful to Dr Mehmet Fisek and Dr Adam M. Packer for initial advice on the optical system and thank Dr David P. Roberson for sharing the NIR-FTIR technology. We gratefully acknowledge feedback on the manuscript from Dr Adam M. Packer and Professor John N. Wood. This work was support by a Sir Henry Dale Fellowship jointly funded by the Wellcome Trust and the Royal Society (109372/Z/15/Z).

