## Supplemental Table and Figures for "Scanned optogenetic control of mammalian somatosensory input to map input-specific behavioral outputs"

**Supplementary information**

**Figure 1 - table 1. List of components for the assembly of the optical system**

| Description | Part Reference Number | Notes | Number |
|---|---|---|---|
| **Stimulation** | | | |
| Laser, 473 nm 100 mW | 06-01 MLD, Cobolt | Mounted on heatsink | 1 |
| Laser heatsink | Custom | | 1 |
| Alignment mirrors | BB1-E02, Thorlabs | *M1* and *M2* in Fig. 1 | 2 |
| Alignment mirror mounts | POLARIS-K1-H, Thorlabs | Mounts BB1-E02 mirrors (*M1* & *M2*) | 2 |
| Lens, 30 mm focal length | AC254-030-A, Thorlabs | *L1* in Fig. 1 | 1 |
| Zoom housing | SM1NR1, Thorlabs | Housing for AC254-030-A (*L1*) | 1 |
| Filter wheel | SCFW6, Thorlabs | Houses ND filters (six settings), *F* in Fig. 1 | 1 |
| ND filter, OD 0.5 | NE505B, Thorlabs | Housed in SCFW6 for 32% transmittance (setting 2) or with one NE510B for 3.2% transmittance (setting 5) | |
| ND filter, OD 1.0 | NE510B, Thorlabs | Housed in SCFW6 for 10% transmittance (setting 3) or with one NE505B for 3.2% transmittance (setting 5) | 2 |
| ND filter, OD 1.3 | NE513B, Thorlabs | Housed in SCFW6 for 5% transmittance (setting 4) | 1 |
| ND filter, OD 2.0 | NE520B, Thorlabs | Housed in SCFW6 for 1% transmittance (setting 6) | 1 |
| Lens, 150 mm focal length | AC254-150-A-ML, Thorlabs | *L2* in Fig. 1 | 1 |
| Lens, 500 mm focal length | AC254-500-A-ML | *L3* in Fig. 1 | 1 |
| Lens mount | LMR1/M, Thorlabs | Mounts C254-150-A-ML (*L2*) and AC254-500-A-ML (*L3*) | 1 |
| Dichroic Mirror | FF776-Di01-25x36, Semrock | *DM* in Fig. 1 | 1 |
| Dichroic mirror mount | CM1-DCH/M, Thorlabs | Houses FF776-Di01-25x36 (*DM*) | 1 |
| End cap | SM1CP2, Thorlabs | Blocks one port of CM1-DCH/M | 1 |
| Mirror galvanometer system | GVSM002/M, Thorlabs | x-axis and y-axis mirror galvanometers (*GM*), mount, servos and power supply | 1 |
| Galvanometer mount | GCM002/M, Thorlabs | Houses GVSM002/M mirrors (*GM*) | 1 |
| Translation stage | DT12XYZ/M, Thorlabs | Mounts GCM002/M | 1 |
| Glass platform | Custom cut borosilicate, 230 x 230 x 10 mm | Mice are studied directly on glass platform | 1 |
| Mouse chambers | Custom, laser-cut acrylic | | |
| **Acquisition** | | | |
| LEDs for FTIR | 12v 850 nm LEDs SMD3528-300-IR, 4.8W/m | Surrounds glass platform within aluminium frame | 1 |
| Camera, local high-speed NIR | acA2000-165umNIR, Basler | Acquires at 160 x 160 pixels, 1000 fps | 1 |
| Camera lens | LM12JCM, Kowa | *CL2* in Fig. 1. Attached to acA2000-165umNIR | 1 |

| | | | |
|---|---|---|---|
| Camera, widefield | acA1920-40um, Basler | Acquires at 160 x 160 pixels, 40 fps | 1 |
| Camera lens | LM6HC, Kowa | *CL1* in Fig. 1. Attached to acA1920-40um | 1 |
| LED power supply | 12V DC power | Powers LEDs for FTIR | 1 |
| **Mounting components** | | | |
| Aluminium breadboard base | MB3045/M, Thorlabs | | 1 |
| Post, 38 mm | RS1.5P/M, Thorlabs | Mounts DT12XYZ/M | 1 |
| Post holders, 30 mm | PH30/M, Thorlabs | Holders for TR30/M | 3 |
| Post holders, 30 mm | PH30E/M, Thorlabs | Holders for TR30/M | 4 |
| Posts, 30 mm | TR30/M, Thorlabs | Mounts POLARIS-K1-H (*M1*), CP02/M, SCFW6 (*F*), LMR1/M (*L2* & *L3*), CM1-DCH/M (*DM*) and camera mounts | 7 |
| Post holder, 20 mm | PH20/M, Thorlabs | Holders for TR20/M, mounted on RC1 | 1 |
| Post, 20 mm | TR20/M, Thorlabs | Mounts POLARIS-K1-H (*M2*) | 1 |
| Clamping forks | CF125C/M, Thorlabs | | 3 |
| Zoom housing mount | CP02/M, Thorlabs | Mount for SM1NR1 | 1 |
| Rail | RLA075/M, Thorlabs | Mounts RC1 | 1 |
| Rail carrier | RC1, Thorlabs | Mounts alignment mirror (*M2*) on 20 mm post | 1 |
| Mounting bases | BA2/M, Thorlabs | Mounts acA2000-165umNIR and CM1-DCH/M | 2 |
| Camera mounts | Custom | Mounts acA2000-165umNIR and acA1920-40um | 2 |
| Cage cube connector | CM1-CC, Thorlabs | Connects GCM002/M to CM1-DCH/M | 1 |
| **Control** | | | |
| Master computer | Custom | Controls system and acquires data | 1 |
| Multifunction I/O device | USB-6211, National Instruments | Interface between acquisition computer, galvanometers, Arduino UNO and laser | 1 |
| Arduino | Arduino UNO | Triggers USB-6211 if required | 1 |
| Analysis computer | Custom | Runs scripts and algorithms for quantitative behavioural analysis | 1 |
| **Frame components** | | | |
| Frame rails | 200 x 25 x 25 mm aluminium extrusion | | 4 |
| Frame rails | 375 x 25 x 25 mm aluminium extrusion | | 2 |
| Frame rails | 225 x 25 x 25 mm aluminium extrusion | | 4 |
| Frame rails | 275 x 25 x 25 mm aluminium extrusion | | 6 |
| **Characterization** | | | |
| Power meter | PM121D and S121C, Thorlabs | Measure light intensity at output from mirror galvanometers and at glass platform | 1 |

| | | | |
|---|---|---|---|
| Beam profiler | BP209-VIS/M, Thorlabs | Measure spot size at stimulation plane using alignment plates | 1 |
| Alignment plates | Custom aluminium plate, 275 x 275 x 13.9 mm | Allows accurate placement of BP209-VIS/M at multiple locations at stimulation plane | 1 |
| Grid array | R1L3S3P, Thorlabs | Calibration of galvanometer jump distance at stimulation plane | 1 |

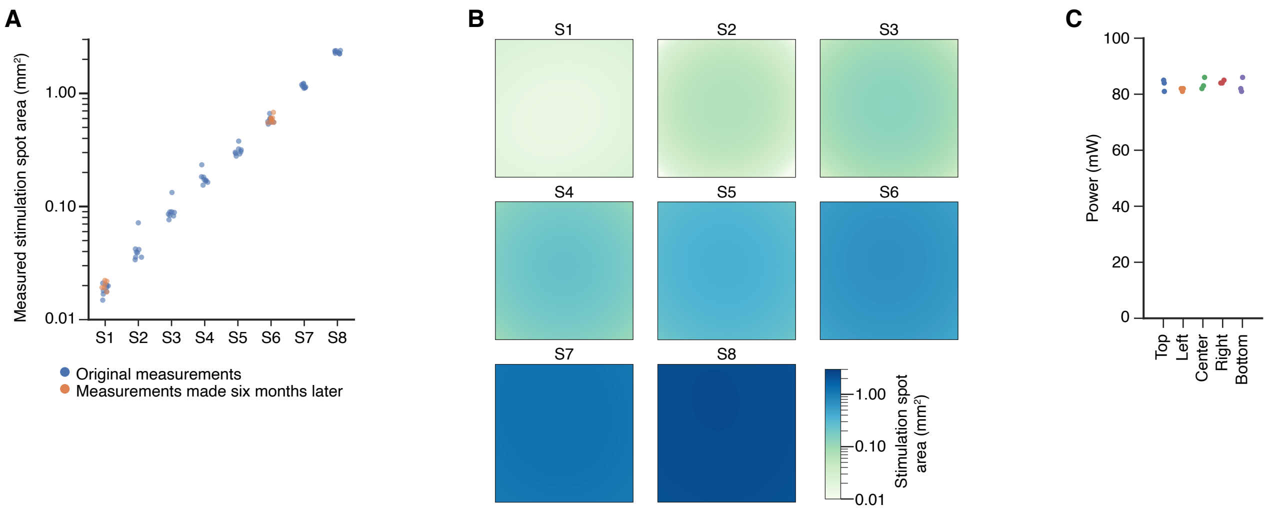

**Figure 1 - figure supplement 1. Technical calibration of optical system.** (**A**) Average spot areas calculated from triplicate measures taken at nine distinct coordinates of the experimental glass platform. The stability of spot size area over time was demonstrated by re-sampling area measurements for S1 and S6 six months after extensive use of the system (orange). (**B**) Uniformity of laser spot area across the surface of the experimental glass platform. Heatmaps of average areas for spot sizes S1 to S8 as measured in triplicates at nine distinct coordinates covering the entire glass platform and fitted with a two-dimensional polynomial equation. (**C**) Uniformity of laser power across the glass platform was demonstrated by measuring laser power in triplicates at five distinct locations using spot size S1 and 100 mW laser output.

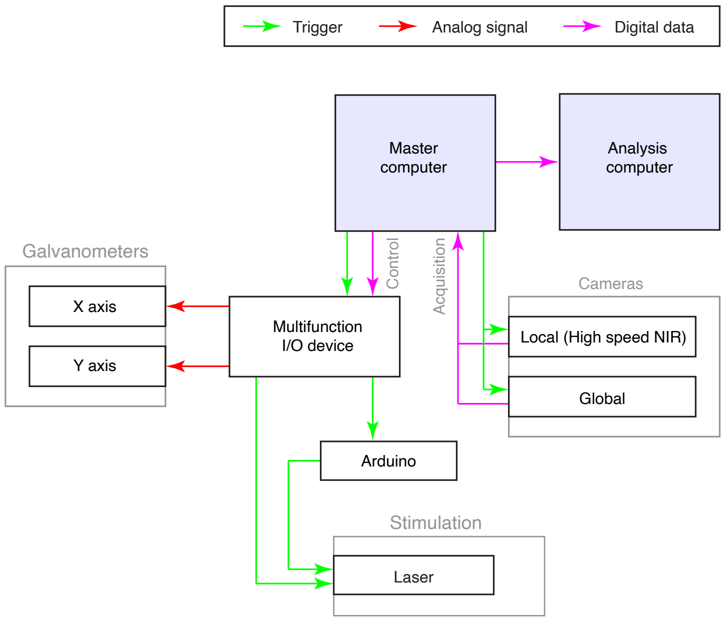

**Figure 1 - figure supplement 2. Hardware and software information flow of the optical stimulation system.** Schematic illustrating the information flow between the system's operating software and the different hardware components. A master computer allows user input to be transformed into digital signals, which are fed into a multifunction I/O device to coordinate the triggering of the laser, cameras and analog control of the galvanometers. The same computer is used to record high-speed paw and full-body behaviors acquired through two separate cameras. Automated analysis is performed offline.

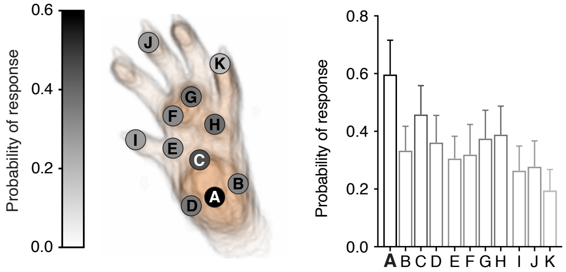

**Figure 2 - figure supplement 1. Microscale mapping of sensitivity to noxious optogentic stimulation.** Paw response probabilities at 11 discrete 0.0185 $mm^2$ stimulation locations acrossthe hind paw glabrous skin using single pulse stimulations (3 ms) in n=8 mice. Response probabilities were determined manually.

**Figure 2 - video 1 (separate file). Pain-related hind paw withdrawals.** Millisecond-timescale changes in hind paw NIR-FTIR signal in response to a single 1 ms laser pulse (laser spot size S5 = 0.577 mm2) recorded at 1000 frames/s. Six individual trials from two different mice are shown.

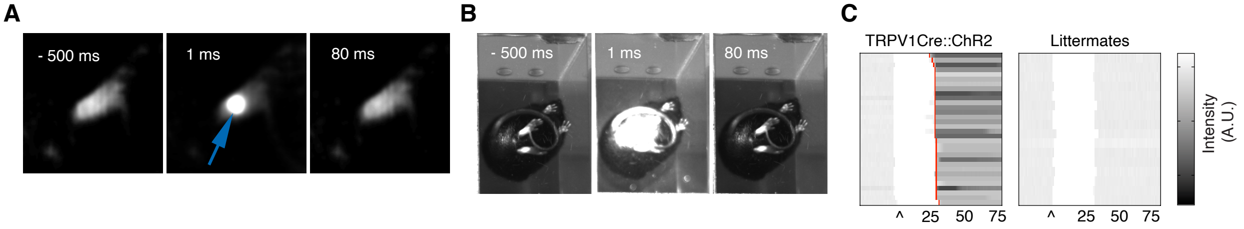

**Figure 2 - figure supplement 2. Littermate controls do not respond to optogenetic stimulation.**
(**A**) Examples of the FTIR hind paw signal before, during and after laser stimulation (arrow).
(**B**) Examples of bottom-view camera recordings before, during and after laser stimulation.
(**C**) Raster plots of hind paw dynamics in response to a single 30 ms pulse (spot size S8 = 2.307 $mm^2$) in TRPV1Cre::ChR2 mice (32 trials from 7 mice) and littermate controls (16 trials from 4 mice). Where applicable, the paw response latency is indicated in red.

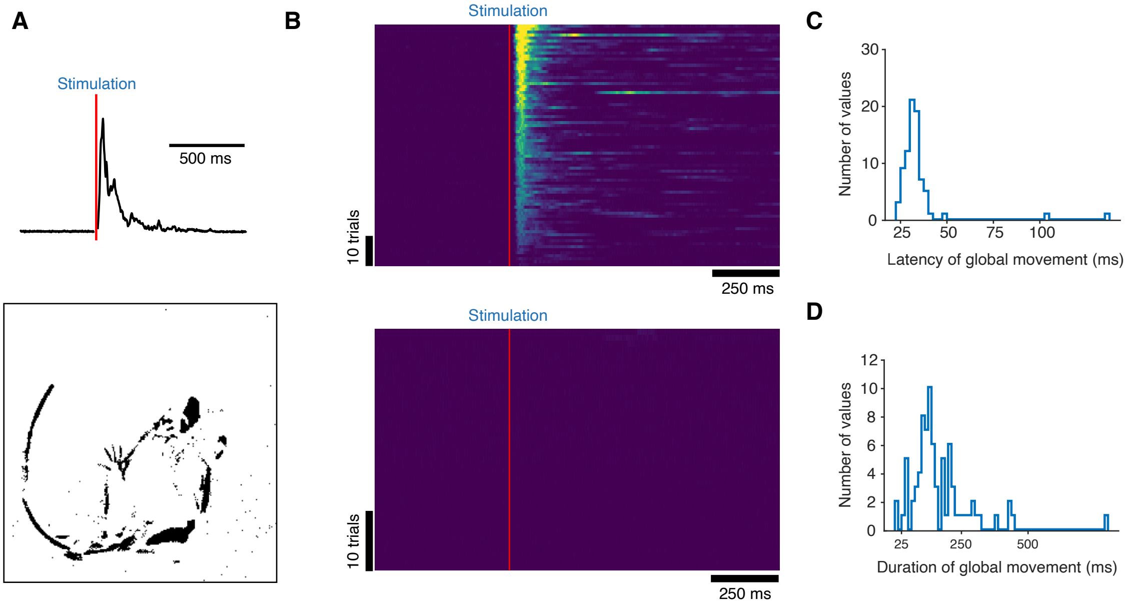

**Figure 4 - figure supplement 1. Motion energy analysis of high-speed recordings.** (**A**) Example motion energy trace acquired from 400 fps videos (top). The stimulus time is shown in red. Example image of animal motion detected by subtraction of neighboring frames (bottom). Pixels that change intensity are shown in black. (**B**) Raster plots of the motion energy time course from ten TRPV1::ChR2 mice from two litters (top; 80 trials from 10 mice) and control littermate mice from the same two litters (bottom; 40 trials from 5 mice). Trials are sorted according to their maximum peak response. The red vertical line represents stimulus. (**C**) Histogram of latencies for stimulus-evoked full-body movements. Latencies were detected at time points when motion energy pixel counts exceeded 10 times standard deviation of the mean baseline signal. (**D**) Histogram of the duration of stimulus-evoked full-body movements. Termination of movement was detected when motion energy pixel counts returned below 10 times standard deviation of baseline signal.

**Figure 4 - video 1(separate file). Pose tracking of behavior in response to nociceptive stimulation.** Markerless tracking was used to analyze postural adjustments in response to a 10 ms optogenetic noxious stimulation (laser spot size S8 = 2.30 mm2). Body parts are labelled with multicolor points and the hind paws and nose connected with magenta lines. The time shown is relative to the stimulus onset.
